# The receptor binding domain of SARS-CoV-2 spike protein fused with the type IIb *E. coli* heat-labile enterotoxin A subunit as an intranasal booster after mRNA vaccination

**DOI:** 10.1101/2023.07.05.547781

**Authors:** He-Chin Hsieh, Chung-Chu Chen, Pin-Han Chou, Wen-Chun Liu, Suh-Chin Wu

**Affiliations:** Institute of Biotechnology, National Tsing Hua University, Hsinchu 30013, Taiwan; Department of Internal Medicine, MacKay Memorial Hospital, Hsinchu 30071, Taiwan; Teaching Center of Natural Science, Minghsin University of Science and Technology, Hsinchu 30401, Taiwan; Biomedical Translation Research Center, Academia Sinica, Taipei 11529, Taiwan; Department of Medical Science, National Tsing Hua University, Hsinchu 30013, Taiwan; Adimmune Corporation, Taichung 42723, Taiwan

**Keywords:** intranasal, RBD, heat-labile enterotoxin A, booster, mRNA vaccine

## Abstract

The outbreak of SARS-CoV-2 infections had led to the COVID-19 pandemic which has a significant impact on global public health and the economy. The spike (S) protein of SARS-CoV-2 contains the receptor binding domain (RBD) which binds to human angiotensin-converting enzyme 2 receptor. Numerous RBD-based vaccines have been developed and recently focused on the induction of neutralizing antibodies against the immune evasive Omicron BQ.1.1 and XBB.1.5 subvariants. In this preclinical study, we reported the use of a direct fusion of the type IIb *Escherichia coli* heat-labile enterotoxin A subunit with SARS CoV-2 RBD protein (RBD-LTA) as an intranasal vaccine candidate. The results showed that intranasal immunization with the RBD-LTA fusion protein in BALB/c mice elicited potent neutralizing antibodies against the Wuhan-Hu-1 and several SARS-CoV-2 variants as well as the production of IgA antibodies in bronchoalveolar lavage fluids (BALFs). Furthermore, the RBD-LTA fusion protein was used as a second-dose booster after bivalent mRNA vaccination. The results showed that the neutralizing antibody titers elicited by the intranasal RBD-LTA booster were similar to the bivalent mRNA booster, but the RBD-specific IgA titers in sera and BALFs significantly increased. Overall, this preclinical study suggests that the RBD-LTA fusion protein could be a promising candidate as a mucosal booster COVID-19 vaccine.

## 1. INTRODUCTION

The outbreak of severe acute respiratory syndrome coronavirus 2 (SARS-CoV-2) in 2019 had led to the coronavirus infection diseases 2019 (COVID-19) pandemic, which has had a significant impact on global public health and the economy (1, 2). SARS-CoV-2 belongs to *Betacolonavirus* of the *Coronaviradae* family and has a single-stranded RNA genome with four major viral structural proteins: spike (S), membrane (M), envelope (E), and nucleocapsid (N) proteins (2–4).The S protein of SARS-CoV-2 is trimeric and contains the receptor binding domain (RBD), which binds to human angiotensin-converting enzyme 2 (hACE2) receptors in human cells (5–12). Numerous RBD-based COVID-19 vaccines have been developed, including those using different expression platforms and adjuvant systems (13–21). Recent improvements to the RBD-based vaccine approach include the use of RBD dimer of delta and omicron (22), heterologous priming with mRNA vaccines (23), and STING-based adjuvants (24) to generate neutralizing antibodies against the immune evasive Omicron BQ and XBB subvariants (25–27).

Nasal immunization can elicit both systemic and mucosal immune responses where the mucosal immune responses such as IgA-secreting B plasma cells can provide a first-line defense against respiratory infections for airway protection (28, 29). However, nasal delivery of vaccines generally requires the use of adjuvants to generate an effective mucosal immunity (29). The most potent mucosal adjuvants are derived from bacterial products such as cholera toxin (CT) and *Escherichia coli* heat-labile enterotoxin (LT) (29–33). Both CT and LT are a member of the AB-class of bacterial toxins; it is composed of an A subunit with ADP-ribosyltransferase activity and a B subunit that mediates binding to eukaryotic cell surfaces (30). The A subunit of LT has been shown to activate dendritic cells and triggered IgG2a, IgA, and Th17 responses when administered intranasally (34, 35). Direct fusion of toxoid antigen to the A subunit has been reported to induce neutralize antitoxin antibodies for enterotoxigenic *E. col*i vaccine development (36). The direct fusion of the CT A1 subunit with influenza A matrix protein 2 (M2e) and an immunoglobulin binding D domain (DD) (CTA1-M2e-DD) has been shown to confer a broad range of protective immunity against homologous and heterologous influenza A viruses (37–39). We have previously reported that a direct fusion of the type IIb LT A subunit (LTA) with influenza hemagglutinin via intranasal immunization induced both systemic and mucosal immunity against H5N1 infections in mice and chickens (40). In this study, we used the direct fusion of the type IIb LTA with RBD of SARS CoV-2 S protein (RBD-LTA) as an intranasal vaccine candidate and evaluated for its elicitation of systemic and mucosal and systemic immune responses in BALB/c mice and golden hamsters. The study also investigates the use of RBD-LTA as an intranasal booster after bivalent mRNA immunization and its potential for developing a mucosal booster COVID-19 vaccine.

## 2. MATERIALS AND METHODS

### 2.1 Expression, purification, and characterization of RBD and RBD-LTA recombinant proteins

The RBD coding sequence containing an N-terminal gp67 signal peptide, a 9x His tag, and the insect cell codon-optimized RBD at residues 330 to 521 of SARS-CoV-2 S protein (GenBank accession no. MN908947.3) was cloned into a pFastBac plasmid (Invitrogen). For the RBD-LTA fusion protein expression, the sequence of E. coli type IIb heat-labile enterotoxin A subunit (GenBank accession no. P43529) was fused to the C-terminal of the RBD construct with a GS linker (G-G-S-G-G-G-S-G). The Bac-to-Bac System (Invitrogen) was used to obtain recombinant baculoviruses according to the manufacturer’s instructions. For large-scale production, Sf9 cells were incubated in 600 mL Sf-900 SFM II serum-free medium (Invitrogen) at a concentration of 2 × 10^6^ cells/mL, and then infected with a specific recombinant baculovirus at a multiplicity of infection (MOI) of 3. Culture supernatants were collected 72 h post infection, and then the recombinant proteins were purified with nickel-chelated affinity chromatography (Tosoh), dialyzed with phosphate-buffered saline (PBS), and stored at 4°C. The purity and molecular weight were determined by 10% sodium dodecyl-sulfate polyacrylamide gel electrophoresis (SDS-PAGE) gels stained with Coommassie blue (0.1% R250) and western blotting using anti-RBD antibody at 1: 10000 dilution (Genetex).

### 2.2 Binding of RBD and RBD-LTA fusion proteins to hACE2 by enzyme-linked immunosorbent assay (ELISA) assay

The experiments were carried out by using 96-well ELISA plates (Thermo) coated with 0.2 μg/well of RBD and RBD-LTA proteins and incubated overnight at 4°C. A recombinant hemagglutinin (HA) protein of Influenza A H1N1 (A/Texas/05/2009) (Sino Biological, Cat. No.40006-V08H) was used as a negative control. The plates were then washed three times with PBST (0.05% Tween-20 in PBS) and blocked with a blocking buffer (1% BSA in PBS) for 2 hours at room temperate (RT). After blocking, the plates were washed three times with PBST. Serial dilutions of hACE2 (Genescript) were added to the wells and incubated at RT for 1 hour. After three washes with PBST, the plates were incubated with 100 µl anti-hACE2 antibody (Genetex) diluted at 1:10000 in PBST for 1 hour at RT. After three washes with PBST, HRP-conjugated anti-rabbit antibody (Genetex) diluted at 1:10,000 in PBST was added to each well and incubated at RT for 1 hour. After the final wash, the plates were incubated with TMB (3,3’,5,5’-Tetramethylbenzidine) (BioLegend) substrate for 15 minutes, and the reaction was stopped by adding 2N H_2_SO_4_ to the well. The optical density was then measured at 450 nm using an ELISA reader (TECAN).

### 2.3 Mouse and golden hamster immunizations

Groups of female BALB/c mice (6-8 weeks old, 5 or 6 mice per group) and golden hamsters (6-8 weeks old, 6 hamsters per group) were intranasally immunized with three doses of RBD-LTA fusion proteins over a 3-week interval. Before immunization, the mice were anesthetized by isoflurane (Panion & BF Biotech) inhalation, while golden hamsters were anesthetized by intraperitoneal injection of Zoletil/Xylazine (Rompun) (20-40mg/kg Zoletil + 5-10 mg/kg Xylazine) and inhalation of isoflurane. The samples of RBD, RBD-LTA, and RBD + poly (I:C) were administered intranasally in specific doses and dripped equally into each nostril of the animals. Bivalent mRNA vaccines (Wuhan-BA.1 and Wuhan-BA.4/5, Moderna) were also used for booster immunization, provided by Hsinchu Mackay Memorial Hospitals. For bivalent RBD-LTA booster, RBD proteins of Omicron BA.1 (Sino Biological, Cat. No. 40592-V08H122) and BA.4/5 (Genetex, Cat. No. GTX137098-pro) were mixed with RBD-LTA. Serum samples were collected at specified times and inactivated by incubation at 56 °C for 30 min to inactivate complements, and stored at –20 °C. BALF samples were aspirated from the trachea with PBS, and centrifuged to collect the supernatants, which were stored at –20 °C. CLNs and splenocytes were collected, ground and strained with a cell strainer (Falcon) in RPMI medium for analysis.

### 2.4 Determination of IgG and IgA titers in sera and bronchoalveolar lavage fluids (BALFs)

The RBD-specific IgG and IgA antibody titers in sera and BALFs were measured using ELISA assay. The experiments were carried out using 96-well plates (Thermo) coated with RBD protein at a concentration of 2 μg/ml and incubated overnight at 4°C. The plates were then washed three times with PBST and blocked with a blocking buffer (1% BSA in PBS) for 2 hours at RT. After blocking, the plates were washed three times with PBST. Serial dilutions of the serum or BALF samples were added to the wells and incubated at RT for 1 hour. After three washes with PBST, the plates were incubated with HRP-conjugated anti-mouse IgG antibody (Genetex) diluted at 1:30000, HRP-conjugated anti-hamster IgG antibody (abcam) diluted at 1:10000, or HRP-conjugated anti-mouse IgA antibody (Bethyl) diluted at 1:50000 for one hour at RT. After the final wash, the plates were incubated with TMB substrate (BioLegend) for 15 minutes, and the reaction was stopped by adding 2N H_2_SO_4_ to the well. The optical density was then measured at 450 nm using an ELISA reader (TECAN). The total IgA concentrations in BALFs were determined by a commercial ELISA kit (Invitrogen) using the same procedures.

### 2.5 Pseudovirus neutralization assay

To produce the pseudoviruses, a plasmid expressing the full-length S protein of SARS-CoV-2 (Wuhan-Hu-1, Alpha, Beta, Delta, Omicron BA.1, Omicron BA.4/5, Omicron XBB.1.5) was co-transfected into HEK293T cells with packaging and reporter plasmids pCMVΔ8.91 and pLAS2w.FLuc.Ppuro (RNAi Core, Academia Sinica), using TransIT-LT1 transfection reagent (Mirus Bio). The pseudoviruses were then harvested and concentrated at 48 hours after transfection, and their titers were estimated based on the luciferase activity of SARS-CoV2-Spp transduction. To assess the neutralizing activity of serum samples, the samples were serially diluted and incubated with 1000 TU of SARS-CoV-2-pseudotyped lentivirus in DMEM (supplemented with 1% FBS and 100 U/mL P/S) for 1 h at 37°C. The mixture was then inoculated with an equal volume of 10,000 HEK-293T cells stably expressing the ACE2 gene in 96-well plates, which were seeded one day before infection. The culture medium was replaced with fresh complete DMEM (supplemented with 10% FBS, 100 U/mL P/S) at 16 h post-infection and the cells were then continuously cultured for another 48 h before being subjected to a luciferase assay (Promega Bright-GloTM Luciferase Assay System). The percentage of inhibition was calculated as the ratio of the loss of luciferase readout (RLU) in the presence of serum to that of the no serum control. The formula used for the calculation was (RLU Control-RLU Serum)/RLU Control. Neutralization titers (IC-50) were measured as the reciprocal of the serum dilution required to obtain a 50% reduction in RLU compared to a control containing the SARS-CoV-2 S-pseudotyped lentivirus only. Neutralization curves and IC-50 values were analyzed using the GraphPad Prism 5 Software.

### 2.6 Determination of T cell responses

Splenocytes and cells from cervical lymph nodes (CLNs) were isolated from immunized mice of each group, pooled, and the seeded in 96-well plates at a density of 5×10^6^ cells for splenocytes or 1×10^6^ cells for CLNs. The cells were then stimulated by 1 ug/ml RBD protein in the total volume of 250 μl of complete RPMI. After 72 hours of induction, the culture supernatants were collected and analyzed for the concentrations of IFN-γ, IL-5, and IL-17A using a commercial ELISA kit (Biolegend). The ELISA kit measured the concentration of these cytokines by utilizing specific antibodies that bind to the cytokines in the sample, followed by detection with an enzyme-linked secondary antibody to provide information on the Th1 (IFN-γ), Th2 (IL-5), and Th17 (IL-17A) cellular responses generated by the immunized mice.

### 2.7 Golden hamster challenge with SARS-CoV-2 virus

Groups of golden hamsters that had received intranasal immunizations with three doses were subjected to a challenge with the SARS-CoV-2 virus. The immunizations were administered 7 weeks prior to the challenge. The hamsters were challenged with 10^4^ plaque-forming units (PFU) of the SARS-CoV-2 virus (hCoV-19/Taiwan/4/2020). After the challenge, the hamsters were sacrificed at two different time points: 3 days post infection (dpi) and 6 dpi. Lung tissues and nasal washes were collected from the sacrificed hamsters for further analysis. To measure the virus titer in the lung tissues, the collected samples were homogenized in 600 µl of DMEM (Dulbecco’s Modified Eagle Medium) containing 2% fetal bovine serum (FBS) and 1% penicillin/streptomycin. After homogenization, the supernatants were separated from the homogenates through centrifugation at 15,000 rpm for 5 minutes. These supernatants were then analyzed to determine the virus titers. In addition to measuring the virus titers, the lung tissues collected from the challenged hamsters were also prepared for histological examination. The tissues were fixed with 10% neutral buffered formalin for 24 hours and then transferred to 70% ethanol for an additional 72 hours. Following fixation, the tissues were embedded in paraffin and trimmed to a thickness of 5 mm. Finally, the samples were stained with hematoxylin and eosin (H&E) to facilitate histological examination.

### 2.8 Statistical analysis

Statistical tests for multiple comparison were performed for all groups (except for the PBS control) in case of the IgG and IgA ELISA data. The results were analyzed using the nonparametric Kruskal-Wallis test, with corrected Dunn’s multiple comparison test, using GraphPad Prism v6.01. Statistical significance has been expressed as follows: *p < 0.05; **p < 0.01; and ***p < 0.001. Neutralization curves were fitted based on the equation of nonlinear regression log (inhibitor) vs. normalized response–variable slope and the corresponding IC-50 NT values were obtained from the fitting curves using GraphPad Prism v6.01.

## 3 RESULTS

### 3.1 Expression, purification, and characterization of SARS-CoV-2 RBD-LTA fusion proteins

The spike RBD gene of SARS-CoV-2 Wuhan-Hu-1 strain (GenBank accession number MN908947.3) was cloned into the pFlastBac-1 vector with an additional N-terminal sequence containing the gp67 signal peptide and nine-repeated histidine for RBD protein expression. Alternatively, the C-terminal LTA fusion gene (GenBank accession number P43529) with a GS linker (G-G-S-G-G-G-S-G) was includedfor RBD-LTA fusion protein expression (**Fig. 1A**). Recombinant baculoviruses were obtained using Bac-to-Bac System. Culture supernatants of infected Sf9 cells were collected at 96 hr post infection and purified by nickel affinity chromatography. Purified RBD and RBD-LTA proteins were analyzed on 10% SDS-PAGE gels with Coomassie blue staining and western blotting (**Fig. 1B**). The molecular weights of RBD and RBD-LTA proteins were observed to be 22 and 50 kDa, respectively. The binding properties of 0.20 μg RBD protein, 0.45 μg RBD-LTA protein (0.20 μg RBD content), and 0.20 μg HA (a negative control) to hACE2 were determined using ELISA assay, and no significant difference was observed in their binding curves to hACE2 (**Fig. 1C**). This indicates that the fusion of LTA at the C-terminus of RBD did not affect the RBD binding to hACE2 and retained the intact RBD conformation to elicit neutralizing antibodies for vaccine design.

**Figure 1.**
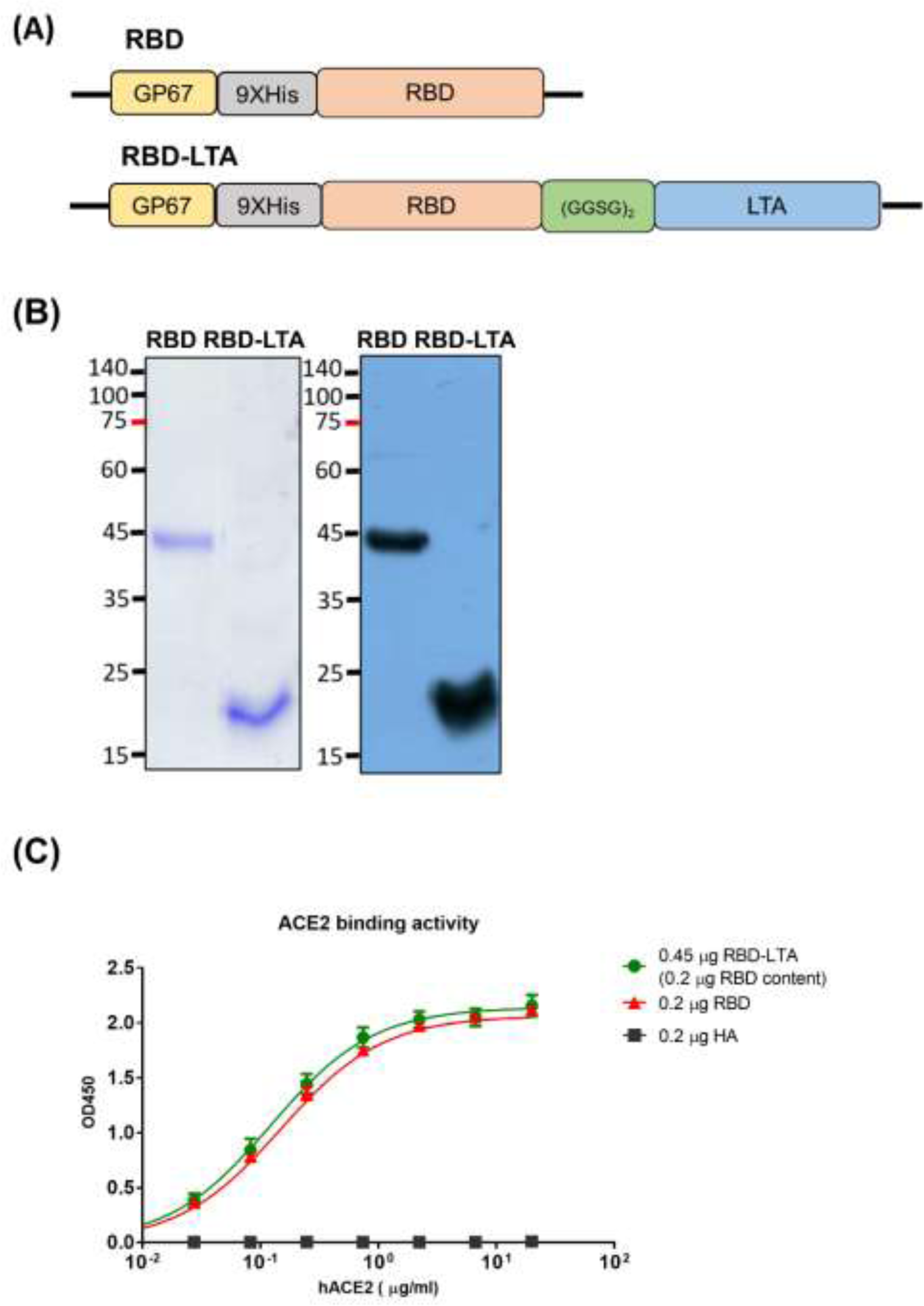
Expression, purification, and characterization of SARS-CoV-2 RBD and RBD-LTA fusion proteins. (A) The expression constructs of RBD and RBD-LTA proteins. **(B)** Purified RBD and RBD-LTA proteins were confirmed by 10% SDS-PAGE gels with Coomassie Blue staining and western blotting. **(C)** ELISA binding curves to hACE2 using 0.2 μg RBD (red), 0.45 μg RBD-LTA (green) with the same RBD content, and 0.2 μg HA (black) as a negative control.

### 3.2 Intranasal immunization with RBD-LTA fusion proteins elicited potent antibody responses in sera and BALFs in BALB/c mice

The immune response to intranasal administration of RBD and RBD-LTA fusion proteins was investigated using RBD + poly (I:C) adjuvant (41) as a positive control. Groups of BALB/c mice (n=5 per group) were intranasally immunized with three doses of (i) PBS, (ii) 20 μg RBD, (iii) 45 μg RBD-LTA (20 μg RBD content), (iv) 20μg RBD +2ug poly(I:C) (**Fig. 2A**). Serum samples were collected two weeks after each dose immunization, and splenocytes, BALFs and CLNs were collected three weeks after the third dose immunization (**Fig. 2A**). Results showed that the RBD-LTA group elicited a significant increase of anti-RBD IgG titer after the third dose intranasal immunization compared to the PBS, RBD, and RBD + poly (I:C) groups (**Fig. 2B**). The RBD-LTA and RBD + poly (I:C) groups elicited higher titers of serum IgA among all immunized groups (**Fig. 2C**). The pooled sera from each immunized group after third dose intranasal immunization were determined for neutralizing antibody titers using pseudovirus assay, and only the RBD-LTA group elicited a significantly IC-50 NT titer against various strains of SARS-CoV-2 **(Fig. 2D).** Furthermore, the RBD-LTA group was found to elicit higher titers of anti-RBD IgA antibodies and higher concentrations of total IgA in BALFs compared to the RBD and RBD + poly (I:C) groups (**Fig. 2E-F**). These results suggest that intranasal immunization with the RBD-LTA fusion protein can elicit both systemic and mucosal immune responses against SARS-CoV-2 infection.

**Figure 2.**
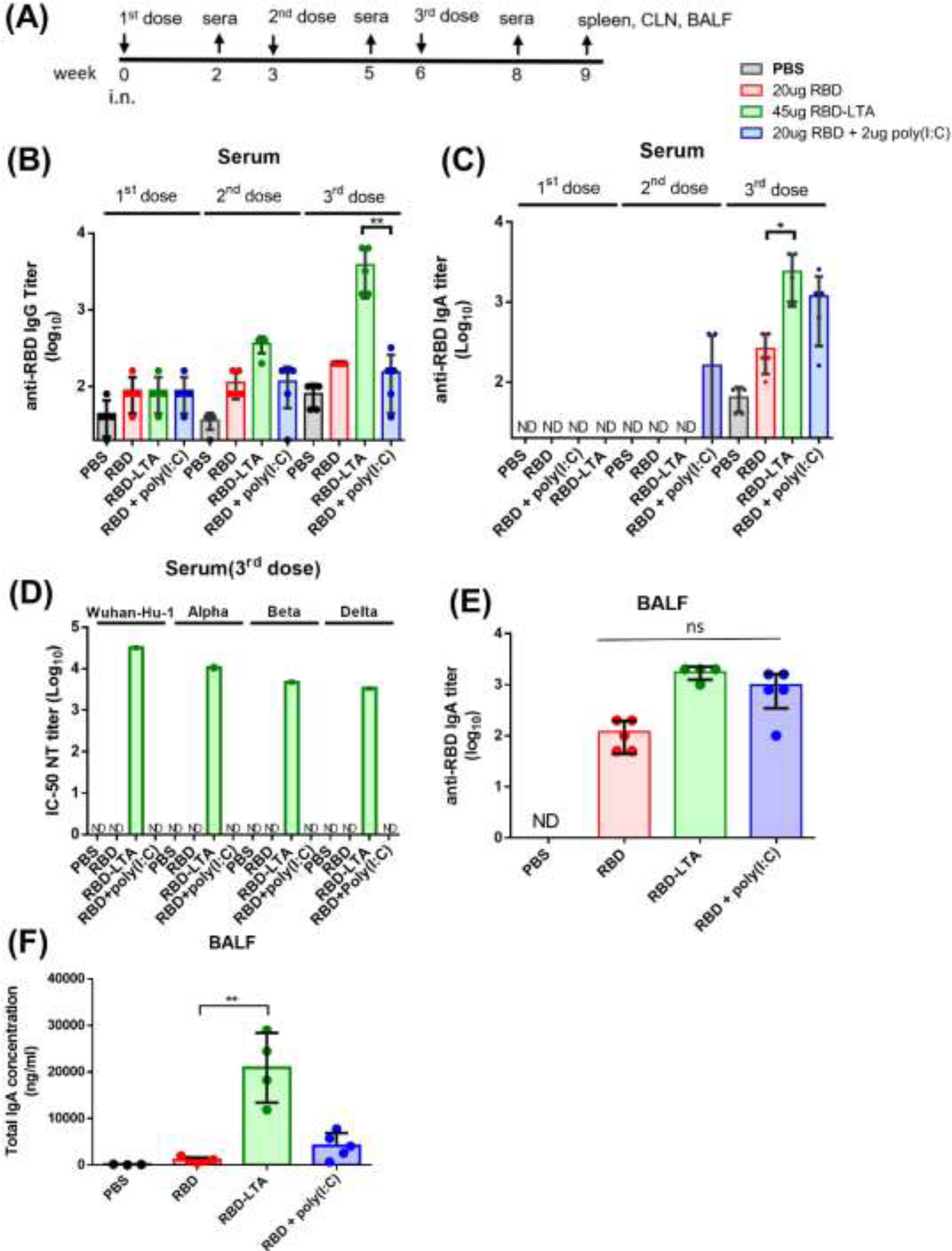
Intranasal immunization with RBD-LTA fusion protein elicited potent antibody responses in sera and BALFs in BALB/c mice. (A) Groups of BALB/c mice were intranasally administered with three doses of PBS, 20μg RBD, 45 μg RBD-LTA, and 20μg RBD + 2μg poly (I:C) in a three weeks interval. Serum was collected 2 weeks after each dose immunization, and all mice were sacrificed three weeks after the third dose immunization to collect splenocyte, CLNs, and BALFs. **(B)** Antisera for anti-RBD IgG titers from each group of mice (n=5) and tested individually after the first, second, and third dose immunization. **(C)** Antisera for anti-RBD IgA titers from each group of mice (n=5) and tested individually after the first, second, and third dose immunization. **(D)** The IC50 values of neutralizing antibody titers after the third dose immunization against Wuhan-Hu-1, Alpha, Beta and Delta. Pooled sera of each immunized group of mice (n=5) and measured in triplicate to obtain dose-dependent neutralization curves using pseudovirus assay. **(E)** The titers of anti-RBD IgA in BALFs from each group of mice (n=5) and tested individually. **(F)** The titers of total IgA in BALFs from each group of mice (n=5) and tested individually. Statistical tests for multiple comparison were performed for all groups (except for the PBS control) in case of the ELISA data. The results were analyzed using the nonparametric Kruskal-Wallis test, with corrected Dunn’s multiple comparison test, using GraphPad Prism v6.01. Statistical significance has been expressed as follows:** p < 0.01. ns: no significance. The IC-50 NT values of neutralization were obtained from the fitting curves of pseudovirus neutralization based on the equation of nonlinear regression log (inhibitor) vs. normalized response – variable slope using GraphPad Prism v6.01. Error bars are plotted as standard deviation from the mean value. Not detectable for N.D.

### 3.3 Intranasal immunization with RBD-LTA fusion proteins elicited Th1, Th2, and Th17 cellular responses in BALB/c mice

We also measured the T cell response to intranasal immunization with RBD-LTA fusion proteins. After three doses of immunization, splenocytes and CLNs of each group were collected, pooled, and stimulated with RBD proteins for 72 h to measure the production of IFN-γ, IL-5, and IL-17A cytokines. The results showed that the RBD-LTA and RBD + poly(I:C) groups elicited higher levels of IFN-γ production compared to the RBD group in spleen and CLNs (**Fig. 3A**). The RBD-LTA group elicited high levels of IL-5 and IL-17A production in spleen and CLNs as compared to all other immunized groups (**Fig. 3C**). Therefore, intranasal immunization with RBD-LTA proteins elicited increased Th1 and Th17 cellular responses.

**Figure 3.**
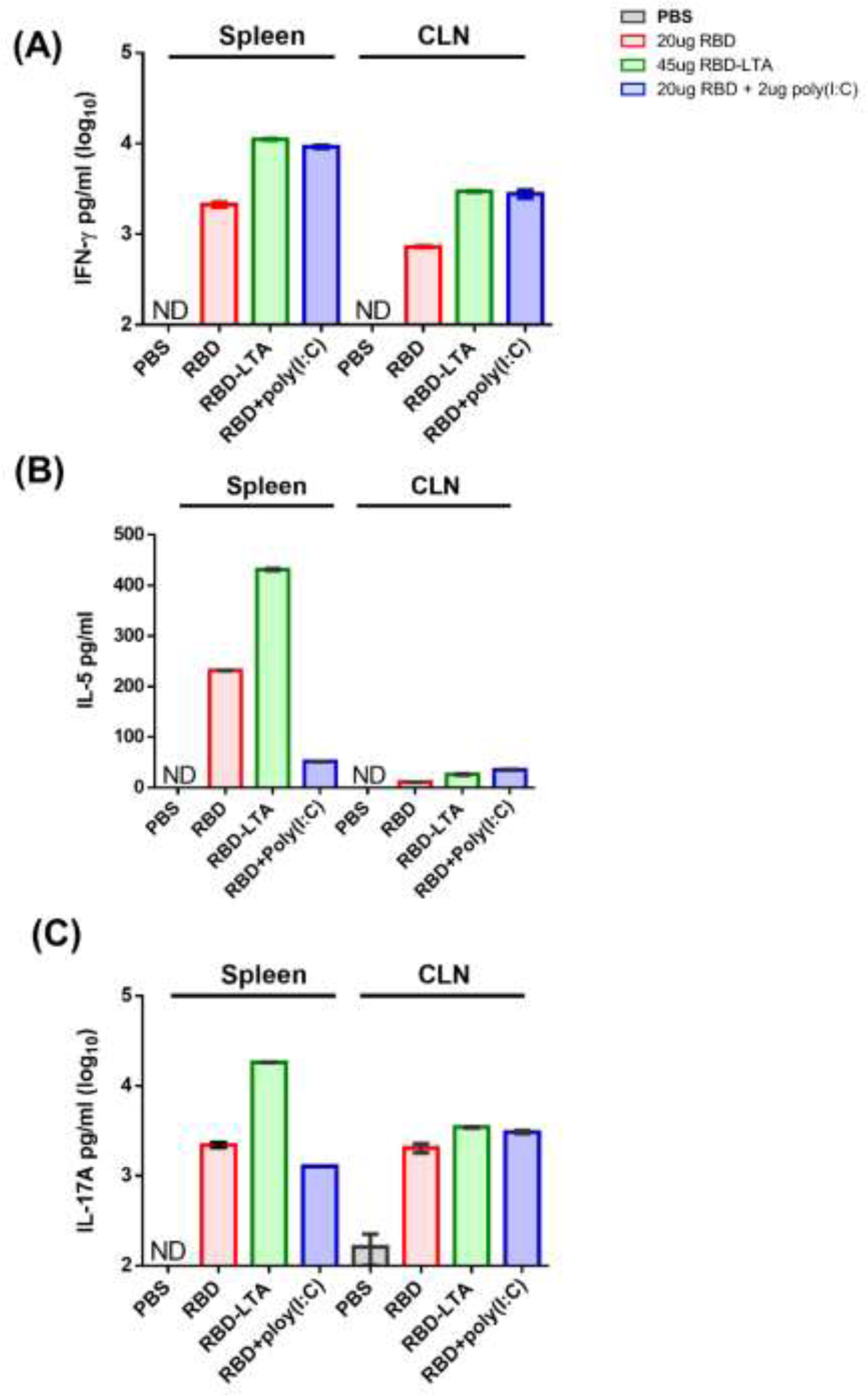
Intranasal immunization with RBD-LTA fusion protein elicited Th1, Th2, and Th17 cellular immunity in BALB/c mice. Splenocytes and CLNs of each immunized group were collected 3 weeks after the third dose immunization, pooled, and stimulated with RBD proteins for 72 h. Culture supernatants were collected and analyzed for **(A)** IFN-γ, **(B)** IL-5 and **(C)** IL-17A cytokine production. Error bars are plotted as standard deviation from the mean value.

### 3.4 Intranasal immunization with RBD-LTA fusion proteins elicited high titers of IgG and neutralizing antibodies and protection in golden hamsters

Golden hamsters were used to investigate the antibody responses elicited by intranasal immunization with RBD-LTA fusion proteins. Three groups of six to eight-week-old hamsters (n=6 pre group) were immunized with PBS, 45 μg RBD-LTA fusion proteins, and 90 μg RBD-LTA fusion proteins, and sera were collected two weeks after each dose of immunization (**Fig. 4A**). The results showed that the groups immunized with 45 μg and 90 μg of RBD-LTA fusion proteins elicited anti-RBD IgG titers reaching approximately 4-5 log scale **(Fig. 4B)** and neutralizing antibody titers around 2-3 log IC-50 values against Wuhan-Hu-01 and the variants of Alpha and Delta but not Beta **(Fig. 4C).** The hamsters immunized with three doses of RBD-LTA and after 7 weeks (i.e. the 13th week after the start of the experiment) were challenged with SARS-CoV-2 by intranasally administering 10^4^ PFU of the virus (hCoV-19/Taiwan/4/2020) (**Fig, 4A**). At 3 dpi, there was no significant reduction in virus titers among all three groups (PBS-immunized, 45 μg RBD-LTA, and 90 μg RBD-LTA) the lung tissue homogenates **(Fig. 4D**). However, at 6 dpi, the group immunized with 90 μg RBD-LTA had the lowest viral titer in the lung tissue homogenates **(Fig. 4D**). At 3 dpi, both the 45 μg or 90 μg RBD-LTA groups had no detectable virus titer in the nasal washes. In contrast, the PBS-immunized group had virus titers ranging from log3 to log4 TCID-50/ml (**Fig. 4E**). The lung tissues from these three immunized groups at 3 dpi and 6 dpi were examined using H&E staining. The results showed moderate lung lesion with the alveolar walls thickened by edema, capillary congestion and variable immune cell infiltration in all three hamster immunized groups at 3 dpi (**Fig. 4F)**. The results from the lungs from the PBS-immunized group at 6 dpi showed a severe endothelial destruction with the shrinking alveoli, hemorrhaging, and mononuclear cell infiltration (**Fig. 4F**). In contrast, the groups of golden hamsters immunized with either 45 μg or 90 μg RBD-LTA retained a moderate inflammatory phenotype with a much less lesion at 6 dpi (**Fig. 4F**)). This suggests that immunization with RBD-LTA fusion proteins provided protection against lung tissue damage caused by SARS-CoV-2 infection. Overall, the results indicate that immunization with the RBD-LTA fusion protein, especially at the higher dose of 90 μg, could effectively reduce virus replication in the lungs, prevent viral replication in the nasal passages, and protect against lung tissue damage caused by SARS-CoV-2 infection in hamsters.

**Figure 4.**
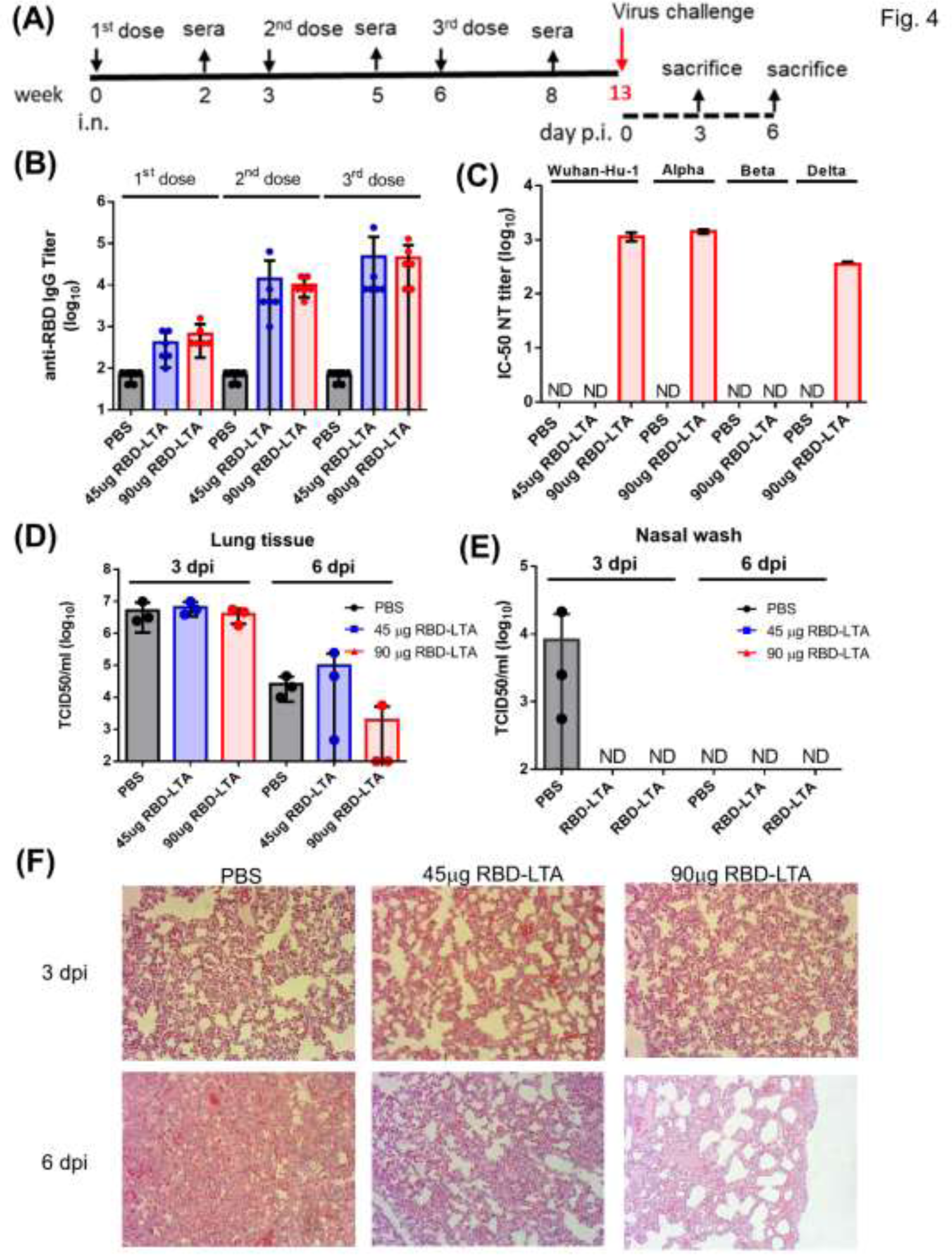
Intranasal immunization of the RBD-LTA proteins in golden hamsters. (A) Groups of golden hamsters were intranasally administered with three doses of PBS, 45 μg RBD-LTA, and 90 μg RBD-LTA in a three weeks interval. Antisera were collected 2 weeks after each immunization. **(B)** Antisera for anti-RBD IgG titers from each group of golden hamsters (n=6) and tested individually after the first, second, and third dose immunization. **(C)** The IC50 values of neutralizing antibody titers after the third dose immunization against Wuhan-Hu-1, Alpha, Beta and Delta. Pooled sera of each immunized group of hamsters (n=6) and measured in triplicate to obtain dose-dependent neutralization curves using pseudovirus assay. **(D)** Virus titers in the lung tissues at 3 dpi and 6 dpi were determined by TCID50 assay. **(E)** Virus titers in the lung tissues at 3 dpi and 6 dpi were determined by TCID50 assay. **(F)** Representative histopathology images of H&E staining from the lung sections of infected hamsters at 3 dpi and 6 dpi, in which purple indicates areas of inflammation. Statistical tests for multiple comparison were performed for all groups (except for the PBS control) in case of the ELISA data. The results were analyzed using the nonparametric Kruskal-Wallis test, with corrected Dunn’s multiple comparison test, using GraphPad Prism v6.01. There are no statistical significances. The IC-50 NT values of neutralization were obtained from the fitting curves of pseudovirus neutralization based on the equation of nonlinear regression log (inhibitor) vs. normalized response – variable slope using GraphPad Prism v6.01. Error bars are plotted as standard deviation from the mean value. Not detectable for N.D.

### 3.5 RBD-LTA fusion proteins as a mucosal booster after bivalent mRNA vaccination in BALB/c mice

The intranasal RBD-LTA fusion proteins were further investigated as a mucosal booster after bivalent mRNA vaccination in BALB/c mice. Two bivalent mRNA vaccines, including (i) Wuhan-Hu-01 + Omicron BA.1 (Moderna) and (ii) Wuhan-Hu-01 + Omicron BA.4/5 (Moderna), were used in the studies. Groups of BALB/c mice (n=6 per group) were first intramuscularly immunized with 2μg bivalent mRNA vaccines, then intranasally boosted with 22.5μg RBD-LTA (10 ug RBD of Wuhan-Hu-01) + 10μg RBD of Omicron BA.1 (**Fig. 5A**) or 22.5μg RBD-LTA (10 μg RBD of Wuhan-Hu-01) + 10μg RBD of Omicron BA.4/5 (**Fig. 6A**). The results were parallel compared with two doses of intramuscular immunization with 2μg/dose bivalent mRNA vaccines. The results indicated that the second dose of the mRNA booster and the intranasal RBD-LTA booster both increased anti-RBD IgG titers against the Wuhan-Hu-01 and Omicron BA.1 (**Fig.5B)** or against the Wuhan-Hu-01 and Omicron BA.4/5 (**Fig. 6B)**. Moreover, the neutralizing antibody titers elicited by intranasal RBD-LTA booster immunization were similar to the bivalent mRNA booster against the Wuhan-Hu-01 and Omicron BA.1 (**Fig. 5C)** or the Wuhan-Hu-01 and Omicron BA.4/5 (**Fig. 6C)**. More interestingly, the intranasal RBD-LTA booster immunization resulted in higher RBD-specific IgA titers in sera (**Figs. 5D, 6D)** and BALF (**Figs. 5E, 6E)**. The total IgA titers in BALFs retained the same levels among all groups against the Wuhan-Hu-01 and Omicron BA.1 (**Fig. 5F)** but significantly increased by the RBD-LTA group against the Wuhan-Hu-01 and Omicron BA.4/5 (**Fig. 6F)**. Therefore, RBD-LTA fusion proteins can be used as a mucosal booster after mRNA vaccine immunizations, and they may offer advantages in terms of increasing mucosal immunity.

**Figure 5.**
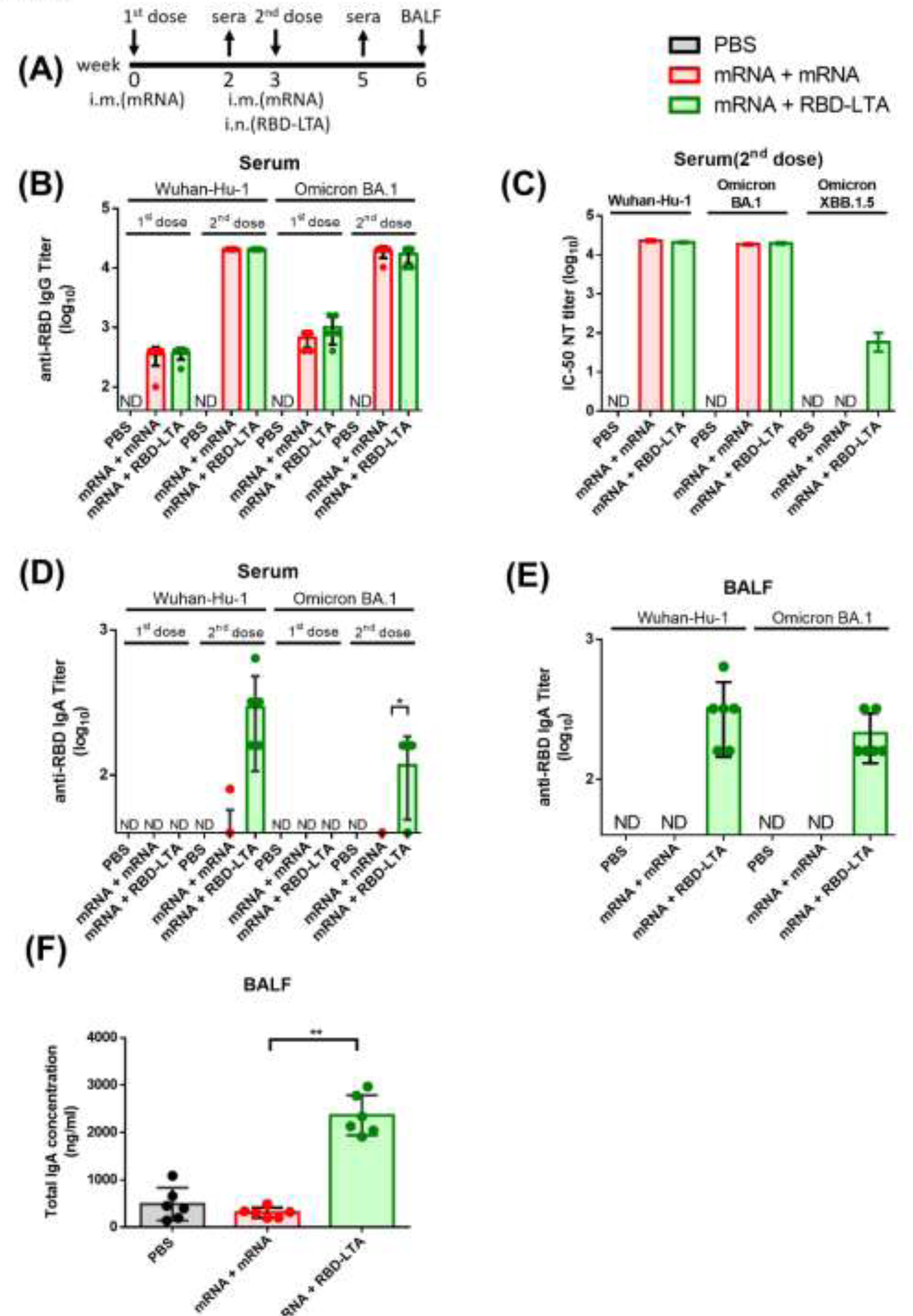
RBD-LTA fusion proteins as a mucosal booster after Omicron BA.1 bivalent mRNA vaccination in BALB/c mice. (A) Groups of BALB/c mice (n= 6) were intramuscularly immunized with either two doses of PBS, 2μg Omicron BA.1 bivalent mRNA, or the first dose of 2μg Omicron BA.1 bivalent mRNA followed by the intranasal booster of 22.5 μg RBD-LTA + 10 ug RBD (BA.1) in a three weeks interval. Antisera were collected 2 weeks after each dose immunization. BALF samples were collected 3 weeks after the second dose immunization. **(B)** Antisera for anti-RBD IgG titers from each group of mice (n=6) and tested individually after the first and second immunization. **(C)** The IC50 values of neutralizing antibody titers after the second dose immunization against Wuhan-Hu-1, Omicron BA.1 and XBB.1.5. Pooled sera of each immunized group of mice (n=6) and measured in triplicate to obtain dose-dependent neutralization curves using pseudovirus assay. **(D)** Antisera for anti-RBD IgA titers from each group of mice (n=6) and tested individually after the first and second immunization. **(E)** The titers of anti-RBD IgA in BALFs from each group of mice (n=5) and tested individually. **(F)** The titers of total IgA in BALFs from each group of mice (n=5) and tested individually. Statistical tests for multiple comparison were performed for all groups (except for the PBS control) in case of the ELISA data. The results were analyzed using the nonparametric Kruskal-Wallis test, with corrected Dunn’s multiple comparison test, using GraphPad Prism v6.01. Statistical significance has been expressed as follows: * p < 0.05, ** p < 0.01. The IC-50 NT values of neutralization were obtained from the fitting curves of pseudovirus neutralization based on the equation of nonlinear regression log (inhibitor) vs. normalized response – variable slope using GraphPad Prism v6.01. Error bars are plotted as standard deviation from the mean value. Not detectable for N.D.

**Figure 6.**
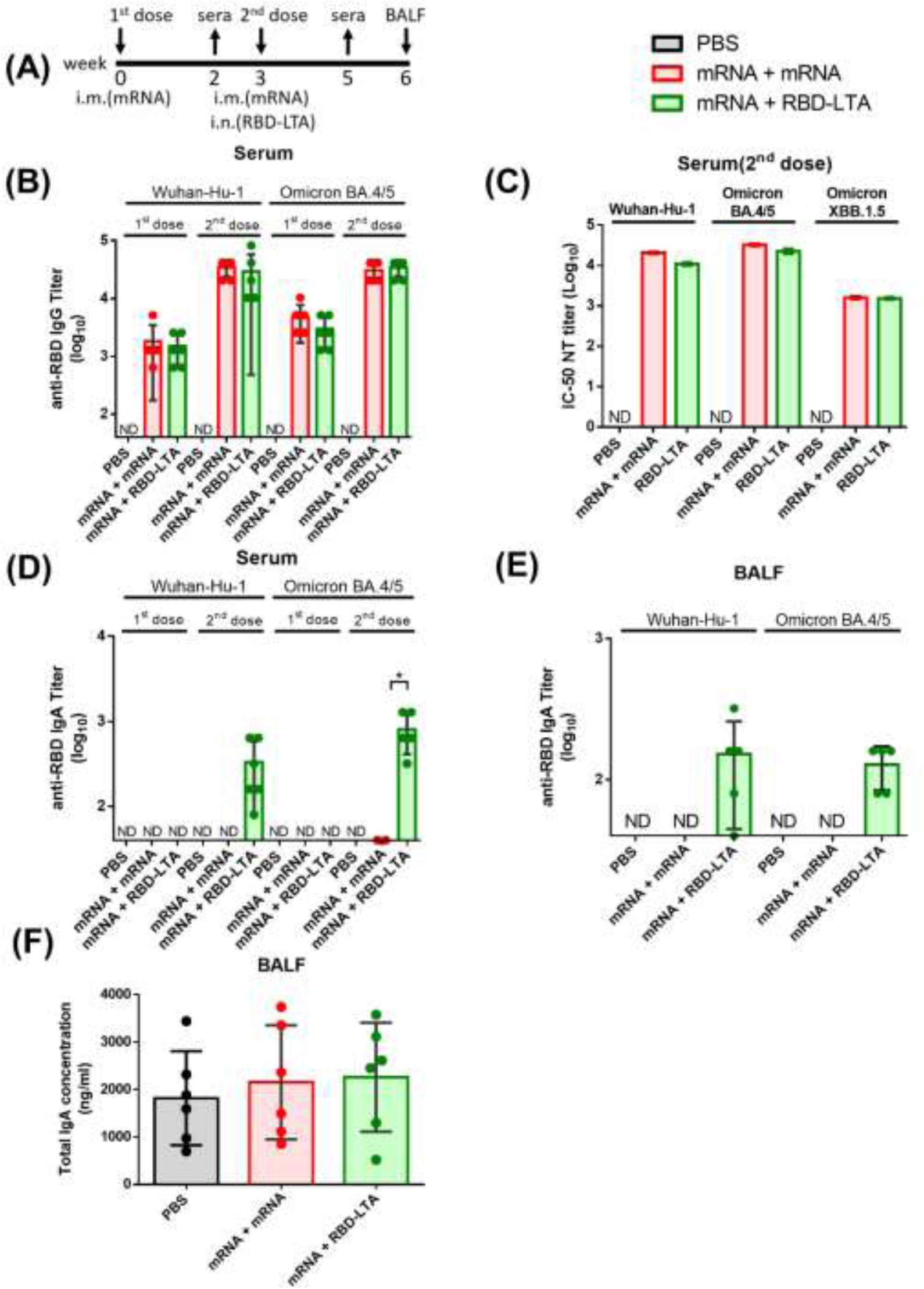
RBD-LTA fusion proteins as a mucosal booster after Omicron BA.4/5 bivalent mRNA vaccination in BALB/c mice. (A) Groups of BALB/c mice (n= 6) were intramuscularly immunized with either two doses of PBS, 2μg Omicron BA.1 bivalent mRNA, or the first dose of 2μg Omicron BA.4/5 bivalent mRNA followed by the intranasal booster of 22.5 μg RBD-LTA + 10 μg RBD (BA.4/5) in a three weeks interval. Antisera were collected 2 weeks after each dose immunization. BALF samples were collected 3 weeks after the second dose immunization. **(B)** Antisera for anti-RBD IgG titers from each group of mice (n=6) and tested individually after the first and second immunization. **(C)** The IC50 values of neutralizing antibody titers after the second dose immunization against Wuhan-Hu-1, Omicron BA.4/5 and XBB.1.5. Pooled sera of each immunized group of mice (n=6) and measured in triplicate to obtain dose-dependent neutralization curves using pseudovirus assay. **(D)** Antisera for anti-RBD IgA titers from each group of mice (n=6) and tested individually after the first and second immunization. **(E)** The titers of anti-RBD IgA in BALFs from each group of mice (n=5) and tested individually. **(F)** The titers of total IgA in BALFs from each group of mice (n=5) and tested individually. Statistical tests for multiple comparison were performed for all groups (except for the PBS control) in case of the ELISA data. The results were analyzed using the nonparametric Kruskal-Wallis test, with corrected Dunn’s multiple comparison test, using GraphPad Prism v6.01. Statistical significance has been expressed as follows:* p < 0.05. The IC-50 NT values of neutralization were obtained from the fitting curves of pseudovirus neutralization based on the equation of nonlinear regression log (inhibitor) vs. normalized response – variable slope using GraphPad Prism v6.01. Error bars are plotted as standard deviation from the mean value. Not detectable for N.D.

## 4 DISCUSSION

It is promising that the use of LT A subunit fusion proteins for developing COVID-19 mucosal vaccines. However, it is important to thoroughly evaluate the safety and efficacy of any potential vaccine candidates. The use of LT B subunit for developing COVID-19 mucosal vaccines has also shown promise in two recent studies (42, 43), but safety concerns must be addressed before the vaccines can be approved for widely use. Previous experience with intranasal influenza vaccine composed of influenza antigens with *E. coli* derived LT-I adjuvant resulted in the total withdrawal of the vaccine trial due to a strong association with Bell’s palsy (44). The mechanism underlying Bell’s palsy after intranasal vaccination remained not totally understood, but the role of LT adjuvants cannot be ignored. The use of LT A-subunit-only adjuvant in vaccines offers the advantage of excluding the concern related to the B subunit binding to GM1 ganglioside of the neuroepithelium (33, 34). This binding to neuroepithelium may have implications for the development of Bell’s palsy, as it could potentially interfere with the normal functioning of the nerve cells. However, the clinical trials using a genetically detoxified mutant A subunit of LT-I (LTK63) along or as a vaccine adjuvant resulted in transient facial palsy (44). Therefore, comprehensive investigation of the safety of LTA fusion proteins as an adjuvant for human vaccines are still needed. It’s important to note that vaccine development and safety are complex processes that involve rigorous testing and evaluation. The use of adjuvants and the consideration of potential side effects are carefully assessed during vaccine development to ensure the overall safety and efficacy of the vaccine.

Here we reported the results of a mucosal vaccine approach in which intranasal immunization with RBD-LTA fusion proteins in BALB/c mice elicited potent neutralizing antibodies against the Wuhan-Hu-1 ancestral strain and the SARS-CoV-2 variants of Alpha, Beta, and Delta (**Fig. 2D**). However, the antisera raised in golden hamsters were not able to neutralize the SARS-CoV-2 Beta variant (**Fig. 4C**), and it may suggest that the RBD-LTA fusion protein was less immunogenic in gold hamsters than BALB/c mice. The present results of immunization experiments comparing the immune response in BALB/c mice and golden hamsters using a RBD-LTA fusion protein. We found that in BALB/c mice, three doses of 45 μg RBD-LTA immunization resulted in a neutralizing antibody titer of approximately 4.5 log10 (**Fig. 2D**). On the other hand, in golden hamsters, three doses of 90 μg RBD-LTA immunization led to a neutralizing antibody titer of approximately 3 log10 (**Fig. 4C**). The results indicate that the neutralizing antibody titers induced in hamsters were less potent than those observed in mice. These findings are in agreement with a recent report that suggested lower titers of neutralizing antibodies were induced by RBD immunization in hamsters compared to mice (45). The discrepancy between the two studies may suggest that the RBD-LTA fusion protein used in this study has altered the antigenic form(s) of RBD, enabling more efficient targeting of neutralizing epitopes in the mucosal system.

To improve the RBD-LTA immunogenicity, the study attempted to replace RBD with S1 (**Fig. S1**), but the purified S1-LTA fusion proteins did not elicit detectable neutralizing antibodies in BALB/c mice using the same intranasal immunization (**Fig. S2**). In the context of the study mentioned earlier, the RBD-LTA fusion protein was used, which presents the RBD in a monomeric form. This monomeric presentation allows the RBD to elicit neutralizing antibodies without relying on the trimeric conformation of the S1 subunit. By bypassing the requirement for the trimeric conformation, the RBD-LTA fusion may be more effective in eliciting neutralizing antibodies compared to the S1 subunit alone. In contrast, the S1-LTA fusion protein, which presents the entire S1 subunit, may not be as effective in eliciting neutralizing antibodies. This is because the S1 subunit requires its conformational structure, including the trimeric form, to properly expose neutralizing epitopes. The monomeric presentation of S1 in the fusion protein may limit its ability to induce a robust neutralizing antibody response. The RBD-LTA fusion protein, by presenting the RBD without the constraint of the trimeric conformation, may be a more efficient strategy for eliciting neutralizing antibodies compared to using the S1 subunit alone.

Based on the results of the study, it was found that intranasal administration of RBD-LTA fusion proteins as a booster after two types of bivalent mRNA vaccination can generate comparable levels of IgG and neutralizing antibodies to those induced by two-dose mRNA vaccination (**Fig. 5B-C**, **Fig. 6B-C**). However, only the intranasal RBD-LTA booster resulted in higher levels of IgA production in BALF (**Fig. 5D-E**, **Fig. 6E-E)**, indicating additional mucosal response beyond two-dose mRNA vaccination. This approach is similarly to a recent report on intranasal S protein booster after primary mRNA vaccination, which also induced IgA production at the respiratory mucosa, boosted systemic immunity, and elicited more broadly neutralizing antibodies against sarbecoviruses (46). The heterologous mRNA priming and subunit protein booster immunizations were shown to increase the breadth of neutralizing antibodies against the SARS-CoV-2 variants in mice (23) and non-human primates (47). Although further investigation is needed to determine whether the RBD-LTA fusion proteins can provide effective transmission-blocking, the current findings can provide the information for developing a mucosal booster COVID-19 vaccine.

## AUTHOR CONTRIBUTIONS

*Project conceptualization, supervision, and administration*: Chung-Chu Chen, Suh-Chin Wu. *Data curation*: He-Chin Hsieh, Pin-Han Chou. *Laboratory investigation*: He-Chin Hsieh, Wen-Chun Liu, Pin-Han Chou. *Data analysis and presentation*: He-Chin Hsieh, Wen-Chun Liu, Pin-Han Chou. *Original draft*: Suh-Chin Wu. *Writing—review and editing*: all co-authors.

## ACKNOWLEDGEMENTS

This work was supported by National Science and Technology Council, Taiwan (NSTC111-2327-B-007-002), National Tsing Hua University (112Q2504E1), and the MacKay Memorial Hospital, Taiwan (MMH-TH-112-04).

## DATA AVAILABILITY STATEMENT

Data available on request from the authors.

## CONFLICT OF INTEREST STATEMENT

The National Tsing Hua University has filed a patent application on the RBD-based COVID-19 mucosal vaccine candidates.

## ETHICS STATEMENT

There is no experimental work with humans included in this article. Procedures involving mice followed the guidelines established by the Laboratory Animal Center of National Tsing Hua University (NTHU). Animal use protocols are inspected and approved by the NTHU Institutional Animal Care and Use Committee (IACUC, Protocol No: NTHU-IACUC-11007H040). All procedures involving the hamster in the study were approved by the Institutional Animal Care and Use Committee (IACUC) and Academia Sinica (Protocol No. BioTReC-110-D-028).

## SUPPLEMENTAL MATERIAL

**Supplementary Figure S1.**
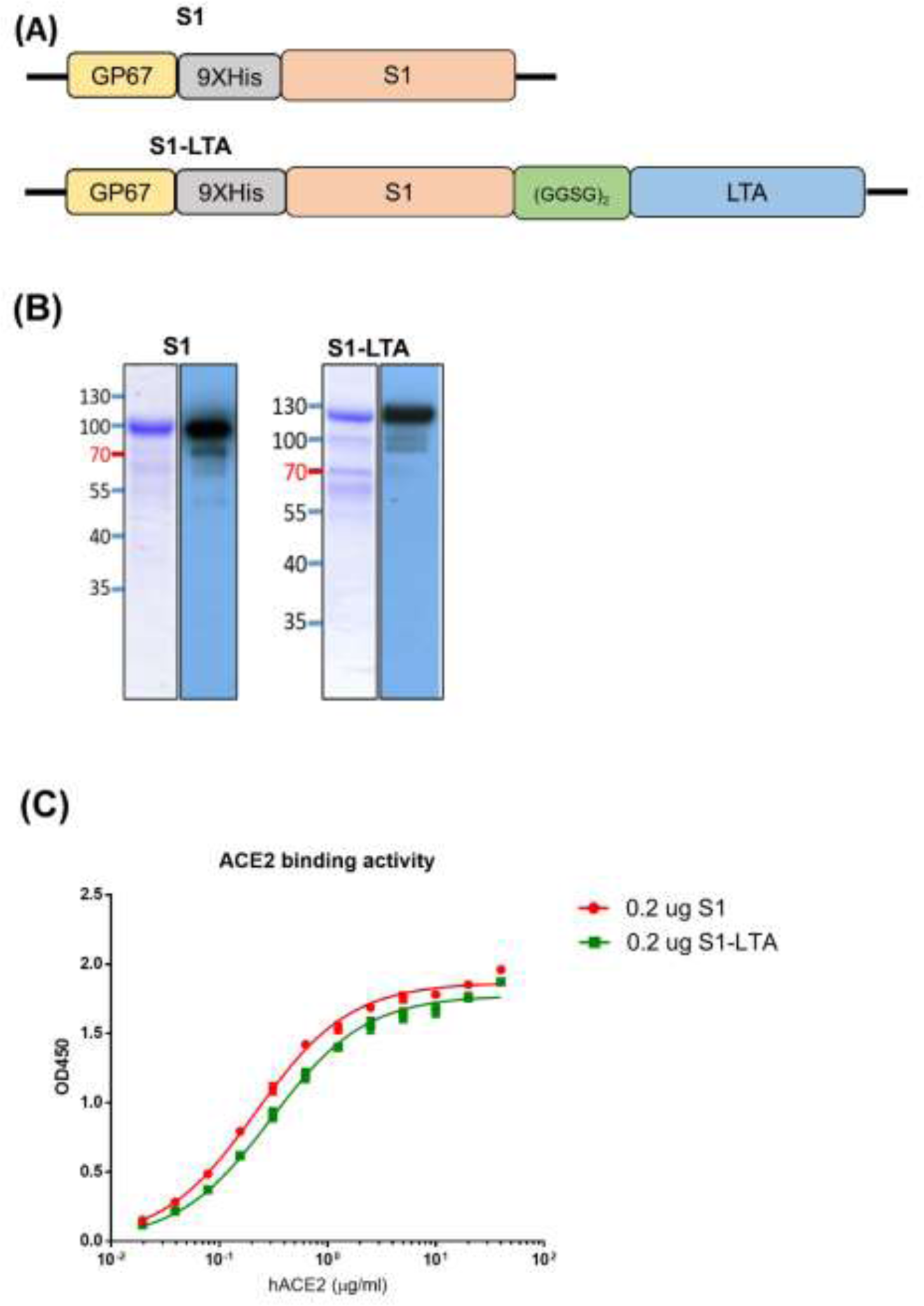
Expression, purification, and characterization of SARS-CoV-2 S1 and S1-LTA fusion proteins. (A) The expression constructs of S1 and S1-LTA proteins. **(B)** Purified S1 and S1-LTA proteins were confirmed by 10% SDS-PAGE gels with Coomassie Blue staining and western blotting. **(C)** ELISA binding curves to hACE2 using 0.2 μg S1 (red) and 0.2 μg S1-LTA (green) with the same RBD content.

**Supplementary Figure S2.**
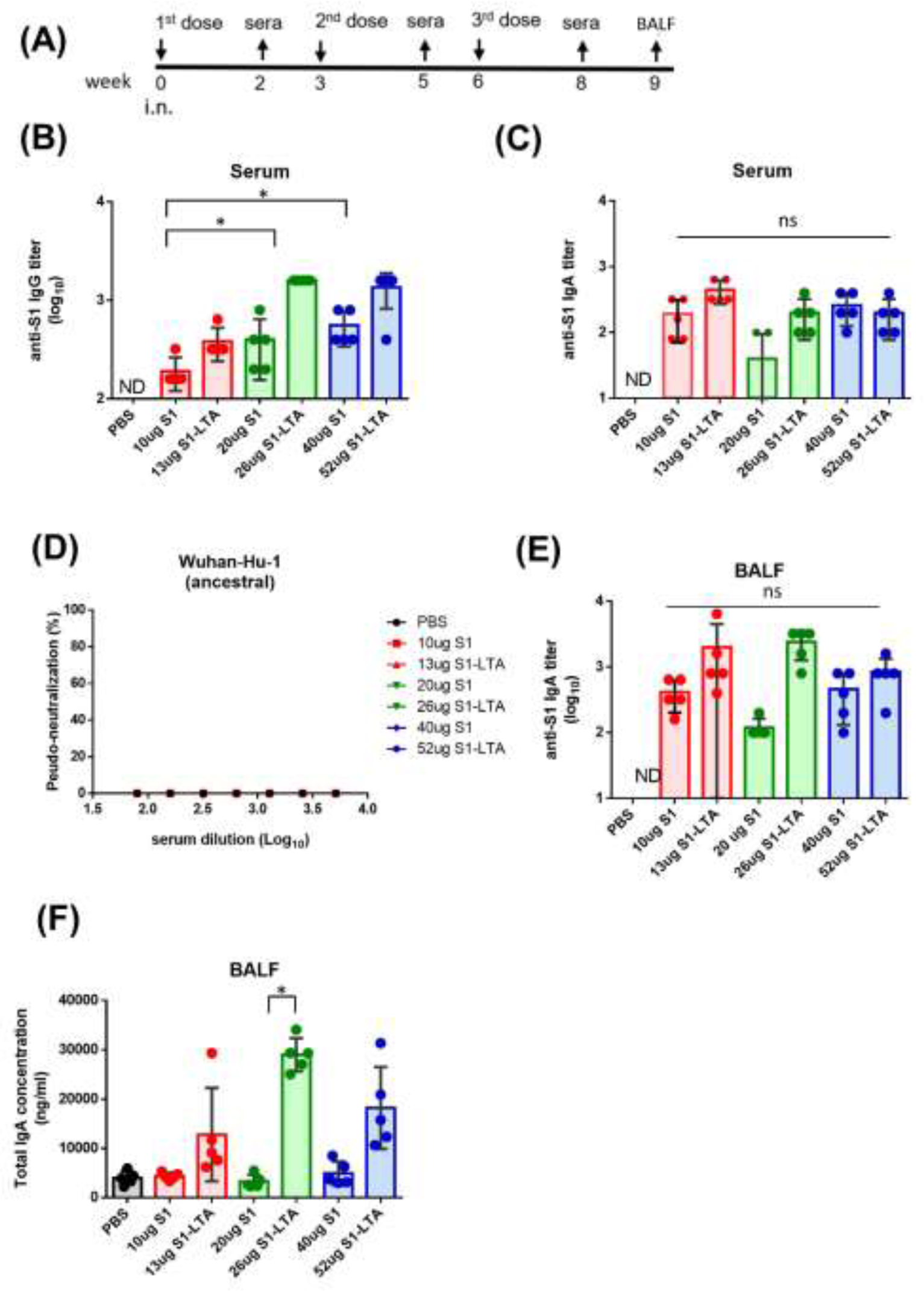
Intranasal immunization with S1-LTA fusion protein elicited potent antibody responses in sera and BALFs in BALB/c mice. (A) Groups of BALB/c mice were intranasally administered with three doses of PBS, 10μg S1, 13μg S1-LTA, 20μg S1, 26μg S1-LTA, 40μg S1, 52μg S1-LTA in a three weeks interval. Serum was collected 2 weeks after each dose immunization, and all mice were sacrificed three weeks after the third dose immunization to collect splenocyte, CLNs, and BALFs. **(B)** Antisera for anti-S1 IgG titers from each group of mice (n=5) and tested individually after the first, second, and third dose immunization. **(C)** Antisera for anti-S1 IgA titers from each group of mice (n=5) and tested individually after the first, second, and third dose immunization. **(D)** The IC50 values of neutralizing antibody titers after the third dose immunization against the Wuhan-Hu-1 strain. Pooled sera of each immunized group of mice (n=5) and measured in triplicate to obtain dose-dependent neutralization curves using pseudovirus assay. The IC-50 values of neutralization were obtained from the fitting curves using GraphPad Prism v6.01. **(E)** The titers of anti-S1 IgA in BALFs from each group of mice (n=5) and tested individually. **(F)** The titers of total IgA in BALFs from each group of mice (n=5) and tested individually. All results were analyzed by GraphPad Prism 6. Statistical significance: * p < 0.05. ns: no significance. Error bars are plotted as standard deviation from the mean value. Not detectable for N.D.

